# High frequency of apomixis in subterranean islands challenges macro-climatic models of geographical parthenogenesis

**DOI:** 10.64898/2026.08.19.745689

**Authors:** Long-Fei Fu, Chi Xiong, Hao Nie, Zi-Bing Xin, Fang Wen, Yi-Gang Wei, Alexandre K. Monro

## Abstract

The Geographical Parthenogenesis (GP) hypothesis traditionally links apomixis to high-latitude and alpine regions, driven by a combination of thermal stress, UV radiation, and pollination barriers. At the core of the GP hypothesis is Baker’s Law, which asserts the evolutionary advantage of autonomous reproduction during the colonization of isolated habitats, frequently linking this advantage to macro-climatic stress. By investigating reproductive modes in subtropical subterranean karst caves, we test the core assumption that macro-climatic stress is the primary driver of apomixis. Our results suggest that barriers to pollination drive a high frequency of apomixis independently of thermal stress or UV radiation. Furthermore, we find a complete phylogenetic turnover between high-elevation, open, apomict-rich habitats and mid-elevation cave environments. Taxonomic composition in these caves is dictated by pre-adaptations to deep shade combined with a lineage-specific predisposition for autonomous reproduction. We propose that the GP hypothesis may be overly influenced by habitat sampling bias. Broader, unbiased sampling could yield a more nuanced and predictive framework to explain variations in apomixis frequency, ultimately providing deeper insights into the role of asexual reproduction in evolution.

## INTRODUCTION

Sexual reproduction in eukaryotes is primarily sustained because of its ability to generate genetic variation and purge deleterious mutations (Crow, 1994; Butlin, 2002; Otto, 2009). Despite this, asexual reproduction occurs across diverse plant lineages (Richards, 1997). Of the several routes to asexual reproduction, apomixis, defined as asexual reproduction via seeds, bypasses meiosis to produce maternal clones (Asker & Jerling, 1992; Bicknell & Koltunow, 2004; Hörandl, 2024). Although apomixis entails long-term genetic costs, such as the accumulation of deleterious mutations through Muller’s ratchet and reduced adaptive potential (Hojsgaard & Hörandl, 2015), it offers immediate ecological advantages by fixing locally adapted genotypes and providing reproductive assurance under pollination limitation, a phenomenon that aligns with Baker’s Law (Baker, 1967). The distribution of apomixis is uneven. Geographically, it exhibits latitudinal and ecological clustering, suggesting environmental drivers. Concurrently, it demonstrates phylogenetic clustering indicative of evolutionary constraints, with approximately 75% of documented apomictic species records concentrated within three major families: Asteraceae, Poaceae, and Rosaceae (Hojsgaard et al., 2014; Hojsgaard & Pullaiah, 2023).

The uneven spatial distribution of asexuality in vascular plants is traditionally explained by the’Geographical Parthenogenesis’ (GP) hypothesis (Vandel, 1928; Hörandl et al., 2008; Hörandl, 2009). The GP hypothesis attributes the disproportionate prevalence of apomicts in high-latitude, high-elevation, and formerly glaciated regions of the Northern Hemisphere (Bierzychudek, 1985; Kearney, 2005) to the colonization advantages of uniparental reproduction in harsh, open, or fragmented post-Pleistocene environments (Hörandl, 2006; Cosendai & Hörandl, 2010; Hörandl, 2011; Schinkel et al., 2016; Tilquin & Kokko, 2016). The GP hypothesis proposes that severe abiotic stressors, such as thermal extremes, UV radiation, shortened growing seasons to pollination, frequently compromise sexual pathways. By bypassing the need for pollination, apomixis provides reproductive assurance, enabling a single colonizing individual to rapidly establish a population while immediately fixing genotypes that are well-adapted to the local conditions (Baker, 1967).

Much of the theoretical consensus surrounding the GP hypothesis is built upon case studies from mountain ranges and extreme alpine or arctic habitats, rather than a systematic sampling of diverse biomes and elevations. In these high-elevation environments, the high frequency of apomixis is conventionally linked to severe abiotic stressors, such as thermal extremes, UV radiation, and barriers to pollination, where it acts as a demographic rescue mechanism under macro-climatic constraints. However, large-scale studies in the NW Himalayas and the American Cordillera have demonstrated that apomixis frequency remains spatially stable regardless of elevational gradients, indicating that broad-scale climatic clines do not consistently drive asexual reproduction (Hörandl et al., 2011; Brožová et al., 2019; Ptáček et al., 2024). This discrepancy suggests the presence of additional ecological filters, such as barriers to pollination, distinct from climate and post-glacial colonization history, that independently drive the transition to asexual reproduction.

Outside of montane and island environments, there has been little systematic study regarding the frequency or drivers of asexual reproduction, for instance, at lower elevations within non-island forest biomes, or concerning its distribution across wider clades. Previous observations of apomixis in the species-rich genus Elatostema (Fu et al., 2017) suggest it may be relatively common in the forest understory and subtropical karst caves, habitats not traditionally associated with this reproductive mode. Subtropical karst caves represent an understudied ecological frontier for plants (Zhu, 2010; Monro et al., 2018; Wei et al., 2025). They contrast sharply with the high-elevation environments central to the GP hypothesis due to their severe light limitation and highly stable, isothermal microclimates. For example, plants in these caves survive at exceptionally low Photosynthetically Active Radiation (PAR) levels of down to 0.35 μmol photons m⁻² s⁻¹ (Monro et al., 2018), ca. 0.02% of full summer daylight (Taiz et al., 2015) and ca. 0.8% to 1.7% of the light available in a deep forest understory (Canham et al., 1990). Southwest China hosts an estimated half a million such caves (Zhang & Zhu, 2021), a significant proportion of which (ca. 30%) harbor vascular plants. The Chinese subtropical karst caves flora comprises ca. 500 species and was primarily assembled from surrounding evergreen broadleaf forests during the Neogene (Monro et al., 2018; Li et al., 2022). Consequently, the highly diverse Chinese cave flora possesses a taxonomic composition fundamentally distinct from the grassland and alpine floras (Asteraceae, Poaceae, Rosaceae) that have dominated previous studies on apomixis (Hörandl et al., 2011; Brožová et al., 2019; Ptáček et al., 2024). Rather, the cave flora is dominated by Gesneriaceae, Urticaceae, and Begoniaceae (Monro et al., 2018; Fu et al., 2022; Wei et al., 2025).

Baker’s Law (Baker, 1955, 1967) describes the distinct evolutionary advantage of autonomous reproduction during the colonization of isolated habitats. Although originally formulated for oceanic islands, it was later applied to high-elevation and extreme environments, which are similarly isolated and fragmented, forming the foundational logic of the GP hypothesis. Our initial observations of apomixis in the Urticaceae (Fu et al., 2017) highlighted the potential for cave colonization via Baker’s Law. Because vegetated karst caves possess discrete boundaries and exceptionally stable microclimates characterized by isothermal temperatures and very low light (Monro et al., 2018), they contrast sharply with high-elevation habitats. This suggested that subterranean caves serve as an ideal natural experiment for testing the robustness of Baker’s Law, specifically isolating the effects of severe barriers to pollination (Li et al., 2006; Wang et al., 2013) from macro-climatic stress.

To leverage this unique ecological setting, we investigated the evolutionary drivers of asexuality within these environments by testing the following hypotheses: 1) that apomixis facilitates the colonization of caves and therefore occurs at a higher rate in cave-dwelling plant populations relative to global averages; 2) that cave apomicts occur predominantly in the same lineages traditionally associated with the GP hypothesis; and 3) if they do not, that cave apomicts are instead drawn from lineages associated with the local forest understory flora that are predisposed to apomixis and low light levels.

## RESULTS

### Overview of Reproductive Mode Screening via B-FCSS

In total, the study employed flow cytometry to analyze 305 populations representing 203 species (Table S2). A subset of samples was excluded from reproductive mode determination for the following reasons: 38 populations (12.46%) lacked a detectable or viable endosperm, while one (0.33%) yielded uninterpretable endosperm-to-embryo ratios. Additionally, in nine populations (2.95%), a secondary peak at the 4C position was observed but excluded from endosperm analysis, as its low nuclei count was consistent with the G2-phase of the embryo rather than a distinct endosperm signal. Consequently, the reproductive mode could be reliably interpreted for 257 populations across 175 species. Of these, 125 species were represented by a single population, while 50 species were represented by two to nine populations. The quality of the flow cytometry data was high, with the mean coefficient of variation (CV) for the embryo tissue peak measuring 3.30% and the mean CV for the endosperm peak being 3.28%.

### Proportion of Apomixis in Subtropical Karst Cave Flora

The study identified four distinct reproductive strategies within the surveyed species pool: sexual reproduction, IRM, autonomous apomixis and pseudogamous apomixis. Sexual reproduction proved to be the overwhelmingly dominant mode, accounting for 81.32% (n = 209) of all observed reproductive events. This was followed by autonomous apomixis, found in 11.28% (n = 29) of the observations, and IRM, present in the remaining 7.00% (n = 18) of the cases and pseudogamous apomixis, found in only 0.39% (n = 1). Further detailed analysis of IRM confirmed a characteristic endosperm-to-embryo ratio in 12 species across six families, with members of the Urticaceae exhibiting this trait most frequently. In contrast, the typical apomictic endosperm-to-embryo ratio was specifically detected in 22 species belonging to 13 genera and 12 families (Table 1).

**Table 1.**
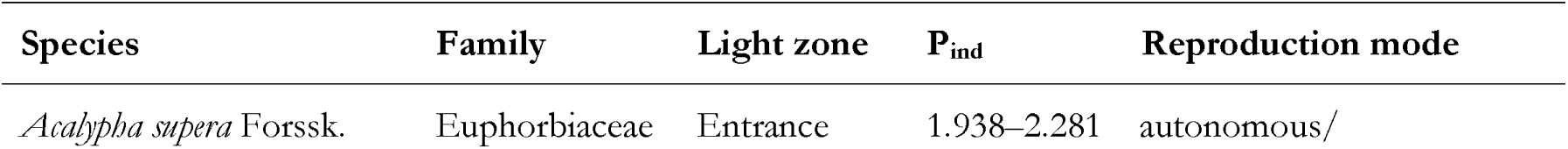
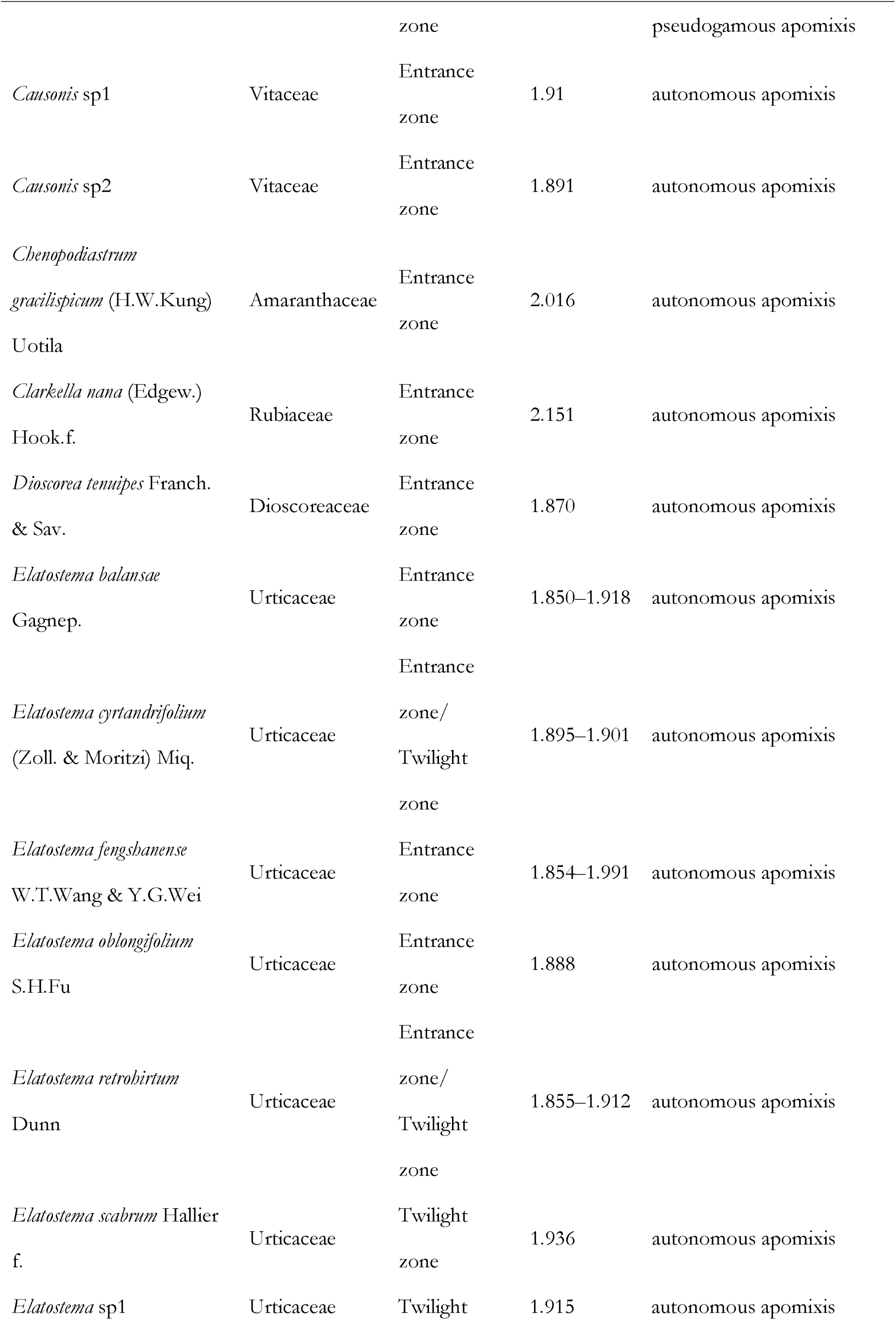
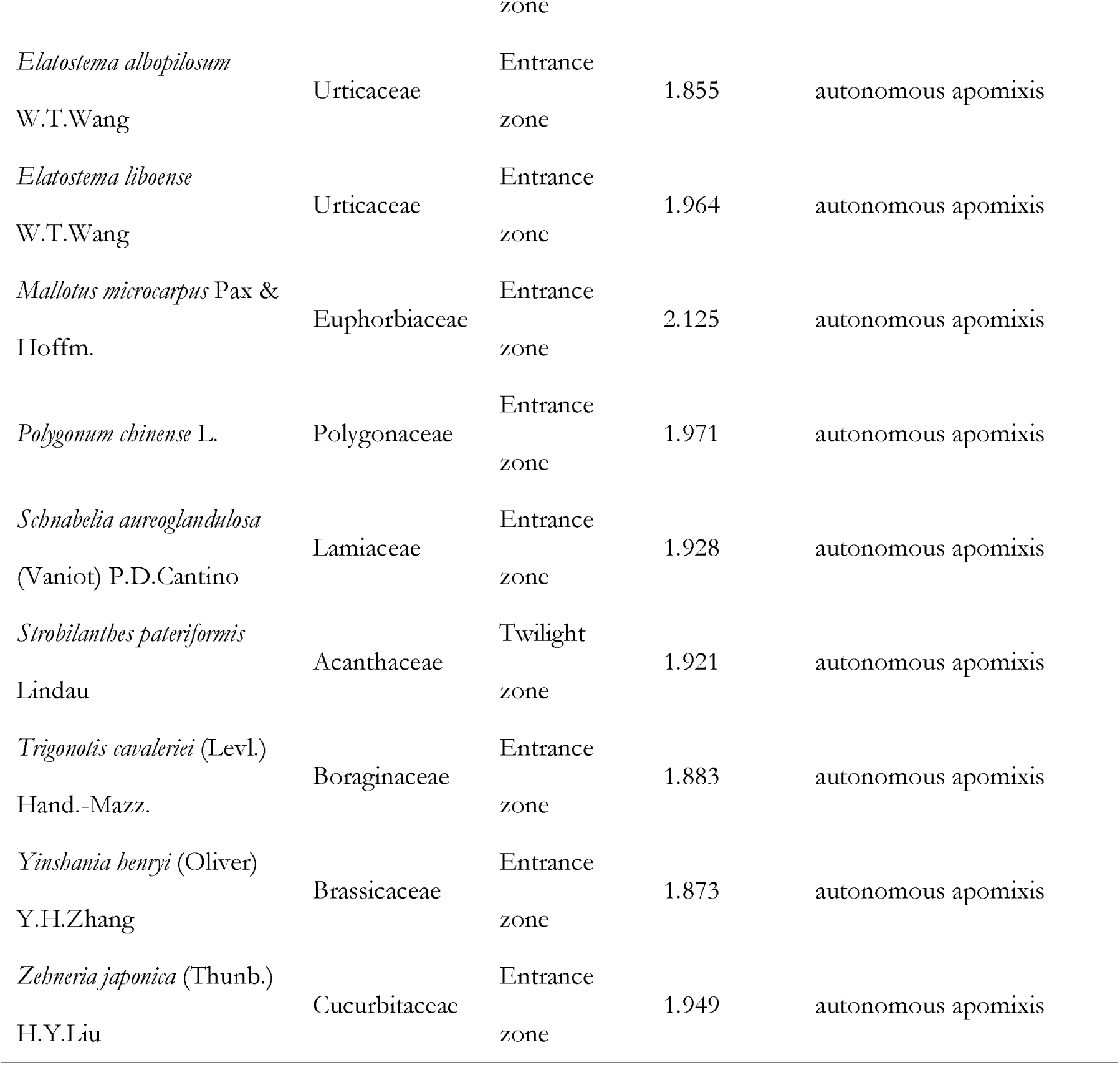
Species list of apomictic species in subtropical karst caves. Endosperm formation in apomictic reproduction is either autonomous (P_ind_ ≈ 2.0) or pseudogamous (P_ind_ ≈ 2.5).

The estimated frequency of apomixis in the subtropical karst cave flora was 12.9% (95% CI: 8.7%–18.7%; 50% CI: 11.2%–14.7%). A conservative sensitivity analysis, conducted to assess the impact of excluding non-endospermic taxa (predominantly 113 species of the cave-dominant Gesneriaceae), revealed an apomixis frequency of 7.7%. While lower than the raw estimate of 12.9%, this conservative frequency is still higher than the Himalayan high-alpine zone (6.0%), and is roughly three times the estimated global angiosperm average of ca. 2.2% (Hojsgaard et al., 2014). When benchmarking this frequency against diverse global alpine floras (Fig. 2), we found that the cave apomixis rate was statistically comparable to those in the northern temperate zone (28.2%; P = 0.083), the southern temperate zone (12.0%; P = 0.965), the tropical alpine zone (15.1%; P = 0.873), and the Himalayan high-alpine zone (6.0%; P = 0.083). The regional proportions of sexual and apomictic species are illustrated in Fig. 2. Within the cave system, the estimated apomixis probabilities in the entrance zone (13.0%; 95% CI: 8.5%–19.4%; 50% CI: 11.2%–15.0%) and the twilight zone (12.0%; 95% CI: 4.2%–30.0%; 50% CI: 8.3%–17.1%) were statistically indistinguishable (P = 0.968), suggesting a uniform distribution of reproductive modes across light gradients (Fig. 2).

**Figure 1.**
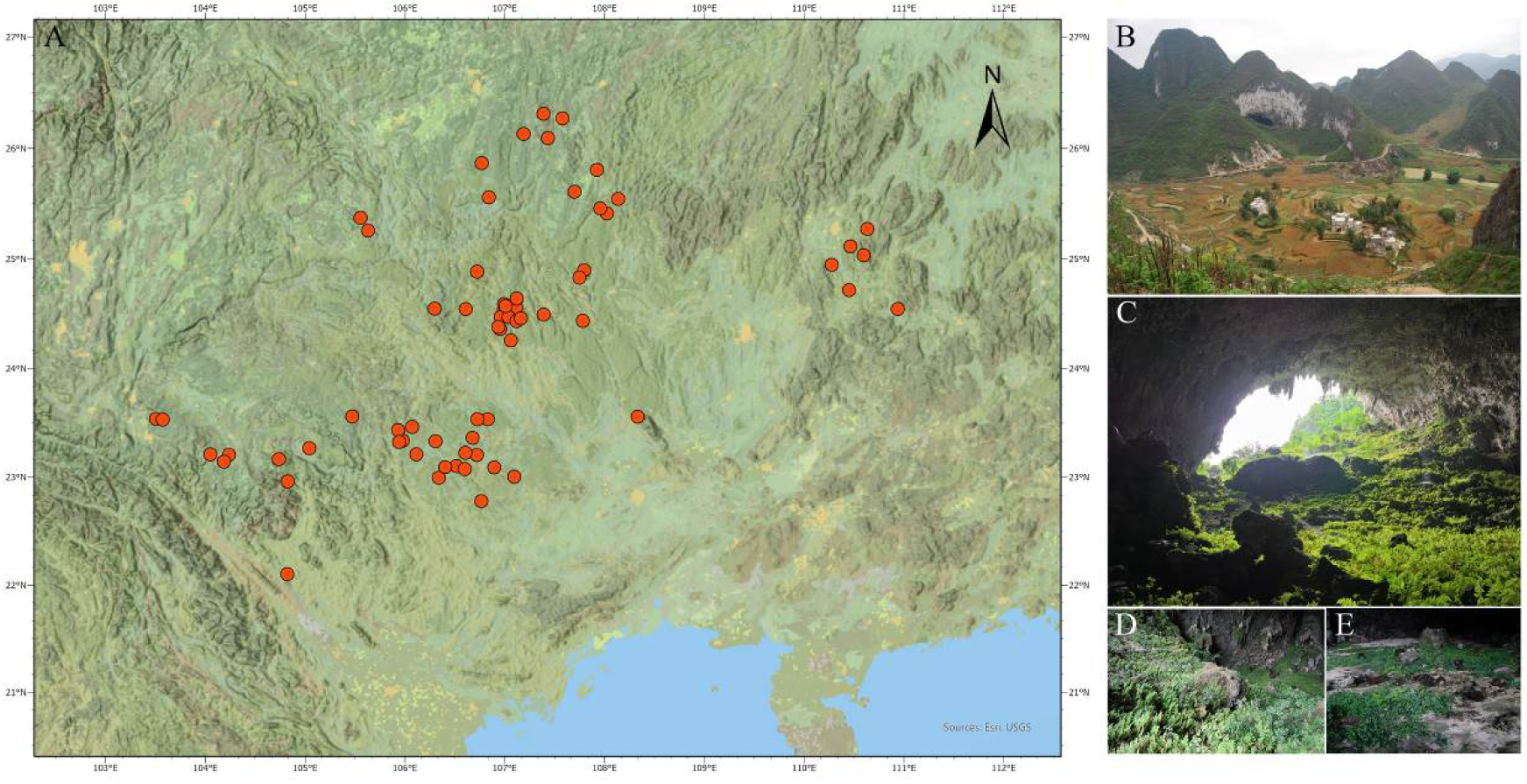
Geographic distribution and habitat characteristics of the sampled cave-associated angiosperms in Southwest China. (A) Topographic map of the study area, where red circles denote the precise locations of the 67 investigated subtropical karst caves. (B–E) Representative field photographs of the sampling environments, depicting (B) an exterior view of the surrounding karst landscape, (C) an interior view looking outward from within a cave, along with specific microhabitats located in (D) the entrance zone and (E) the twilight zone.

**Figure 2.**
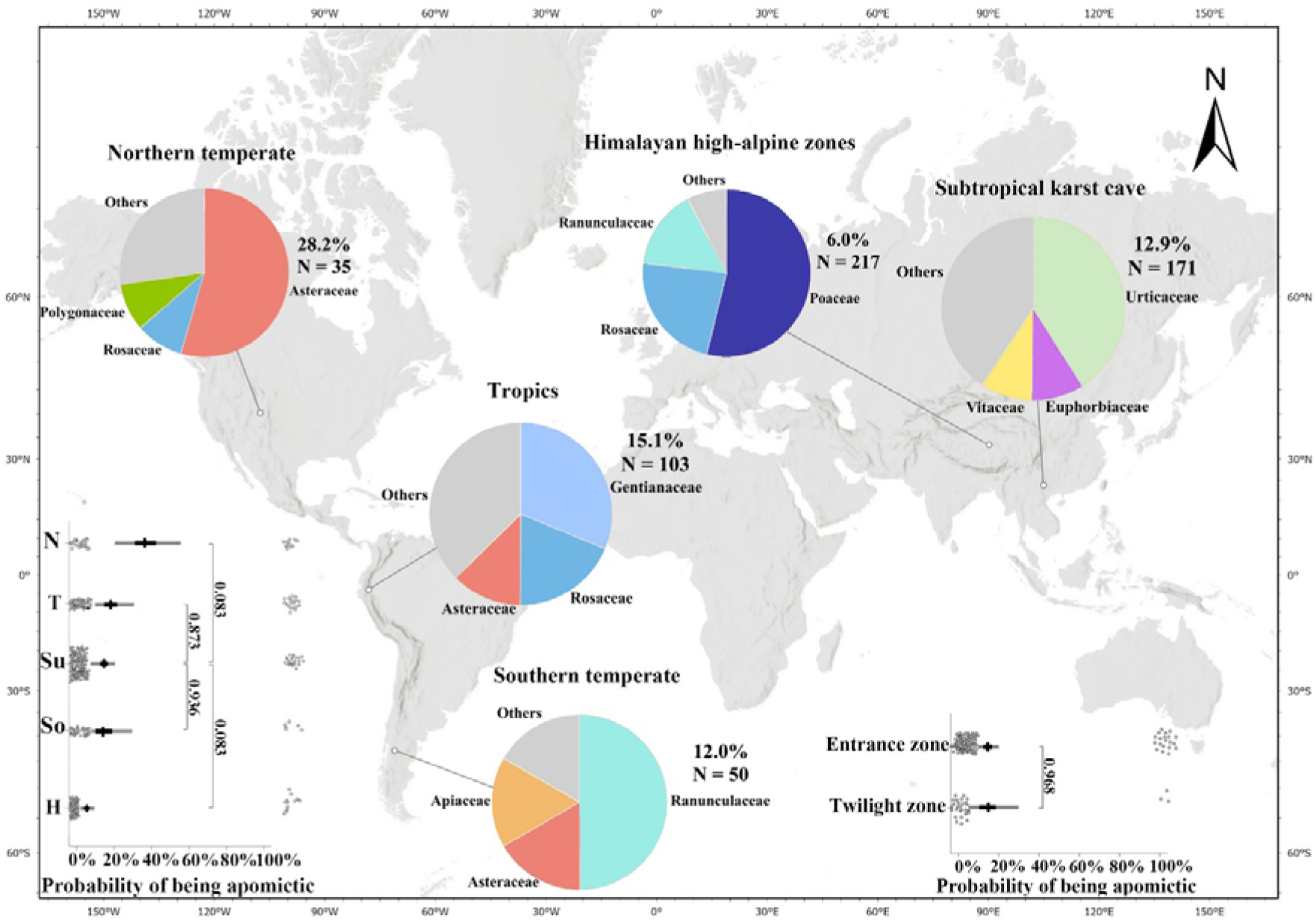
Global context of apomixis prevalence across five biogeographic ecosystems. The world map displays apomictic family distributions across five regions (Northern temperate, Tropics, Subtropical karst cave, Southern temperate, and Himalayan high-alpine). Pie charts show the relative proportions of the top three dominant families in each region’s asexual assembly. Adjacent percentages (%) and N values denote regional apomixis frequency and total sampled species, respectively, directly corresponding to the bottom-left plot. In this plot, points represent mean probabilities, thick/thin bars indicate 50%/95% confidence intervals, and jittered open circles show individual species data. The bottom-right inset compares apomixis probability between the entrance and twilight zones of karst caves. The numerical values adjacent to the brackets in both plots indicate the exact P values for the respective pairwise comparisons.

### Phylogenetic Distribution and Signal of Apomixis in Cave Flora

To determine whether the prevalence of apomixis in cave flora is driven by shared ancestor or independent environmental adaptation, we evaluated the phylogenetic signal across the sampled taxa. The phylogenetic distribution of reproductive modes was mapped onto a family-level tree (Fig. S2). The apomictic reproduction showed a strong phylogenetic signal (estimated s^2^ = 1.153, P = 0.002 for phylogenetic signal equal to zero, Fig. S2). This suggests that apomixis in subtropical karst caves is restricted to a few pre-adapted lineages.

A comparative analysis of phylogenetic turnover was visualized using mirrored trees (Fig. 3). While the Poaceae, Rosaceae, and Asteraceae dominate asexual complexes in high-latitude or alpine regions, these families were either absent or represented only by sexual populations in our cave dataset (Fig. 3; Table 2, S2). Instead, the Urticaceae, Euphorbiaceae and Vitaceae emerged as the primary carriers of apomixis in cave ecosystems, accounting for 40.9%, 9.1% and 9.1% of the identified asexual species, respectively (Fig. 3; Tables 1–2). This shift underscores a unique ecological filtering process where cave-specific environmental stressors favor asexual transitions in lineages with high physiological shade tolerance, effectively filtering out traditional heliophytic ‘asexual-prone’ families.

**Figure 3.**
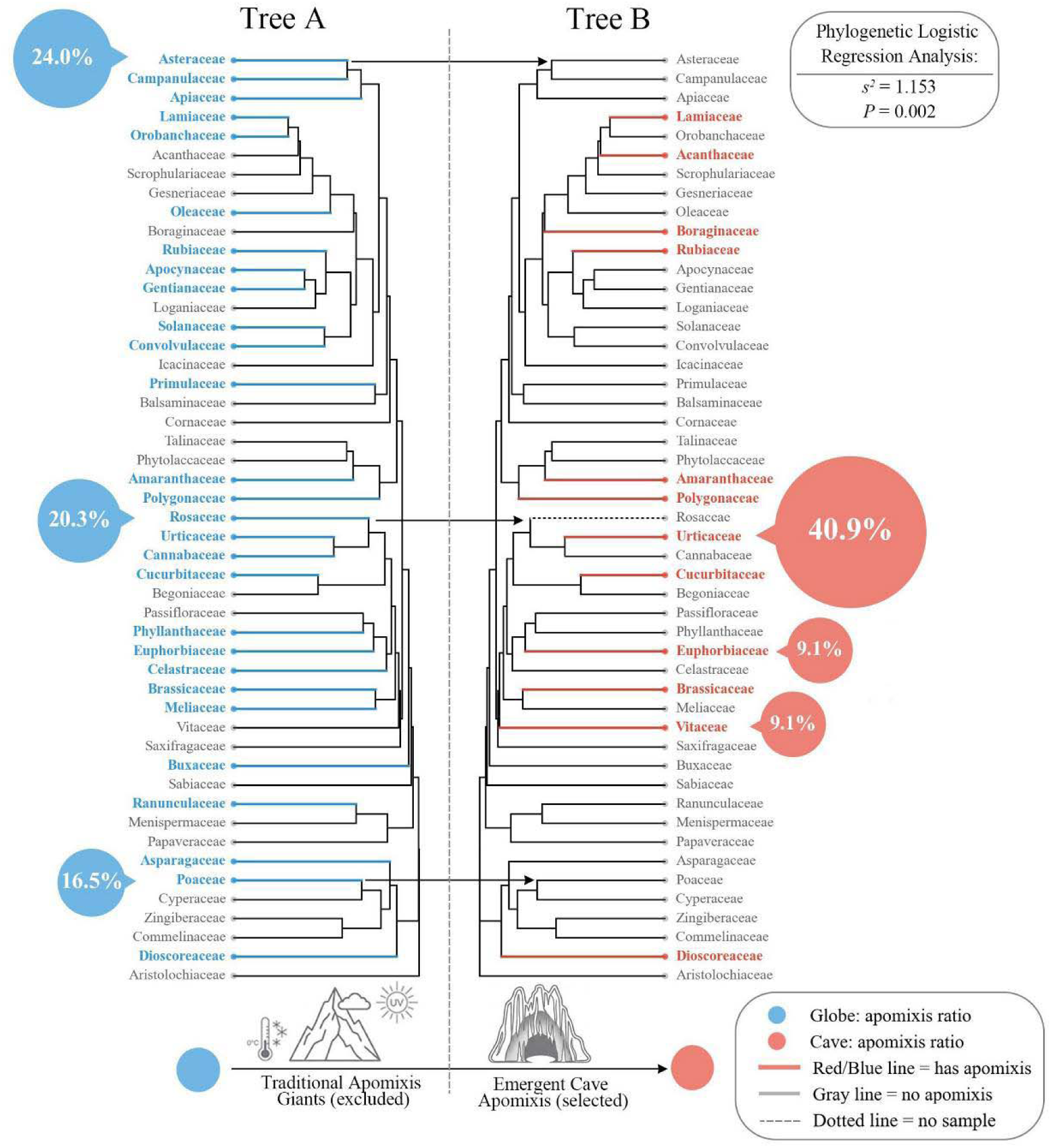
Phylogenetic turnover of apomictic lineages between global conventional ecosystems and subtropical karst caves. The mirrored phylogenetic trees display the identical set of plant families present in the sampled karst cave flora to directly contrast their reproductive behaviors. Tree A (left) illustrates global conventional ecosystems including high-latitude or alpine zones where traditional apomixis giants such as Asteraceae, Rosaceae, and Poaceae dominate asexual reproduction. Tree B (right) demonstrates emergent cave apomixis within the exact same taxonomic pool, showing that these traditional giants reproduce sexually in caves, while apomixis is instead concentrated in shade-tolerant lineages including Urticaceae, Euphorbiaceae, and Vitaceae. Percentages indicate the proportion of total apomictic species accounted for by these dominant families in each respective environment. The upper-right inset displays phylogenetic logistic regression results (s² = 1.153, P = 0.002), confirming a strong phylogenetic signal for reproductive modes in the cave flora.

**Table 2.** Comparative frequency and phylogenetic composition of apomixis across global ecosystems.

| Region/Ecosystem | Dominant Stressor | Apomixis | Dominant | Source |
| --- | --- | --- | --- | --- |
|  |  | Frequency (%) | Families (Apomicts) |  |
| Subtropical Karst Caves | Biotic (Light/Pollinator) | 12.9% (7.7% conservative) | Urticaceae, Euphorbiaceae, Vitaceae | This Study |
| European Alps | Thermal (Cold) | 4.7% | Rosaceae | Hörandl *et al.* (2011) |
| Himalayas (High Alpine) | Thermal (Cold/UV) | 6.0% | Poaceae,<br>Rosaceae | Brožová *et al.*<br>(2019) |
| Northern Temperate | Thermal/Seasonal | 28.2% | Asteraceae | Ptáček *et al.*<br>(2024) |
| Tropical Andes | UV/Diurnal Temp | 12.0% - 15.1% | Asteraceae,<br>Gentianaceae,<br>Rosaceae | Ptáček *et al.*<br>(2024) |

Family composition differed significantly among the five ecosystems in the six-family table and remained significant when all other families were pooled as a seventh category (Fisher-Freeman-Halton tests, both P < 0.001). Thus, the turnover was not an artifact of restricting the comparison to the six focal families.

### Spatial Distribution of Apomixis along Elevational and Latitudinal Gradients

GLMM analysis revealed that apomixis incidence remains statistically stable across the sampled ranges, with neither elevation (122 to 2057 m; Z =-0.669, P = 0.503) and latitude (22°44′ to 26°5′ N; Z = 0.434, P = 0.665) showing a significant correlation with reproductive transitions (Fig. 4). This lack of spatial structure contrasts sharply with open-habitat ecosystems, where reproductive modes typically exhibit predictable responses to climatic or geographic clines. In karst caves, the prevalence of asexual reproduction appears to be a consistent feature of the subterranean environment, independent of external geographic positioning. This suggests that the overarching ecological constraints of caves act as a uniform evolutionary filter for colonizing lineages, effectively overriding the influence of broader macro-environmental gradients.

**Figure 4.**
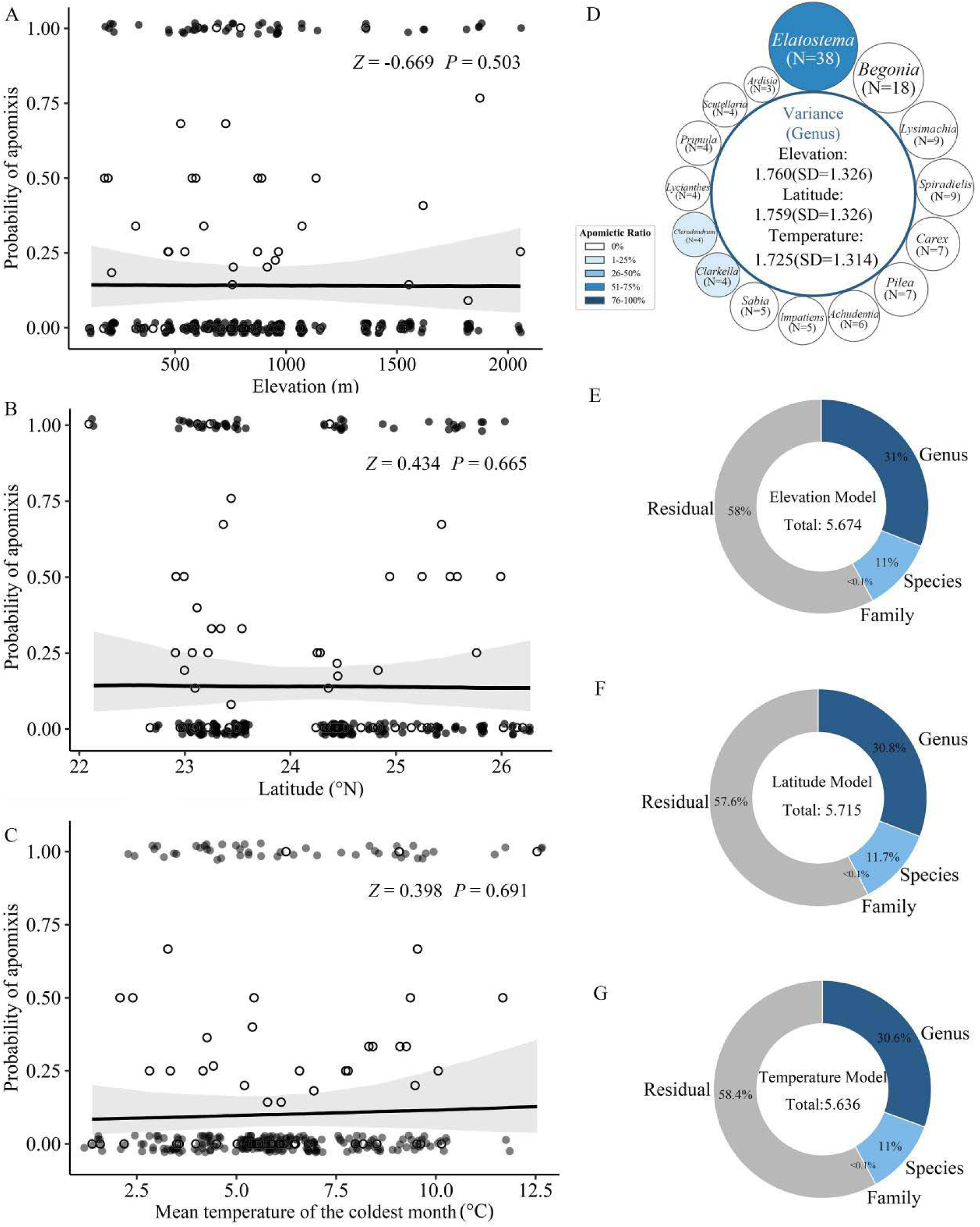
Spatial decoupling of apomixis from macro-climatic gradients and taxonomic variance partitioning. (A–C) Generalized Linear Mixed Models (GLMMs) evaluating the population-level influence of (A) elevation, (B) latitude, and (C) mean temperature of the coldest month on apomixis probability. Solid lines show predicted probabilities with 95% confidence intervals indicated by shaded areas, demonstrating that apomixis frequency remains statistically stable across spatial gradients (P > 0.05). Open circles represent the observed apomixis proportions at each individual cave sampling site. (D) Circular plot detailing variance estimates at the genus level across elevation, latitude, and temperature models, accompanied by the sample sizes (N) and distribution of apomictic ratios within specific genera. (E–G) Donut charts showing variance partitioning across nested taxonomic ranks of Family, Genus, and Species, alongside residuals for (E) the elevation model, (F) the latitude model, and (G) the temperature model.

### Influence of Life Form and Pollination Mode on Apomixis

The GLMM identified life form as a significant predictor of reproductive mode. Annual plants exhibited a higher prevalence of apomixis (35.0%, 7/20) compared to perennial taxa (14.5%, 23/159; β =-1.573, Z =-2.425, P = 0.015; Fig. 6). In contrast, although wind-pollinated species exhibited a higher percentage of apomixis (28.8%, 15/52) than biotic-pollinated taxa (11.8%, 15/127; Fig. 5), this difference was not statistically significant (β =-0.579, Z =-0.993, P = 0.321; Fig. 6). Crucially, the random effect structure across both models revealed that variance was predominantly concentrated at the genus level (Variance: 1.139–1.259), whereas variance at the family and species levels was negligible (Fig. 4).

**Figure 5.**
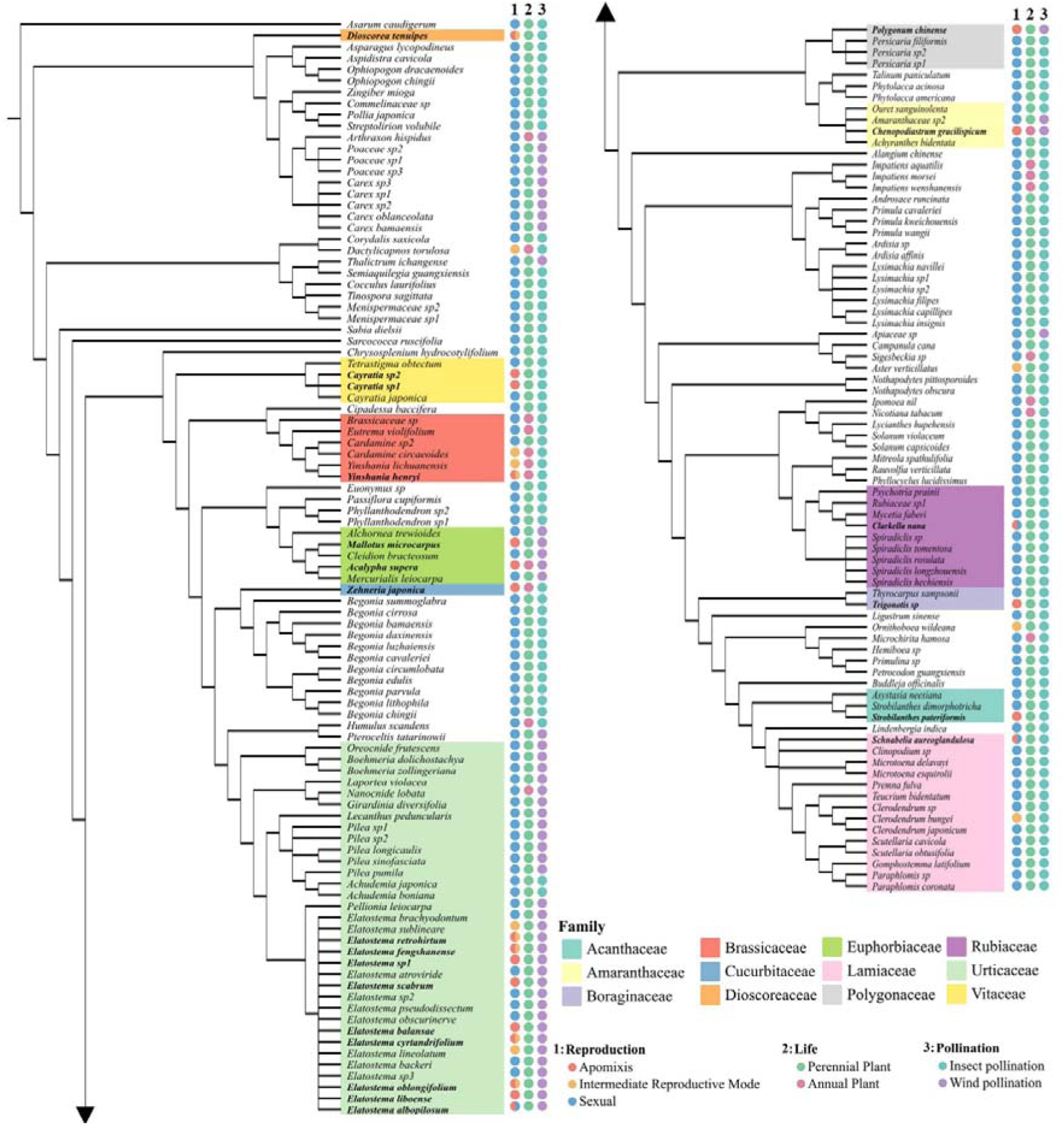
Phylogenetic distribution of reproductive modes and life history traits in subtropical karst caves. A phylogenetic dendrogram mapping angiosperm reproductive strategies, with inner shaded clades highlighting specific plant families. Three adjacent columns display traits corresponding to each tip: (1) Reproduction mode, indicated by orange for apomixis, yellow for intermediate reproductive mode, and blue for sexual reproduction; (2) Life form, indicated by green for perennial plants and pink for annual plants; and (3) Pollination method, indicated by purple for insect pollination and light blue for wind pollination.

**Figure 6.**
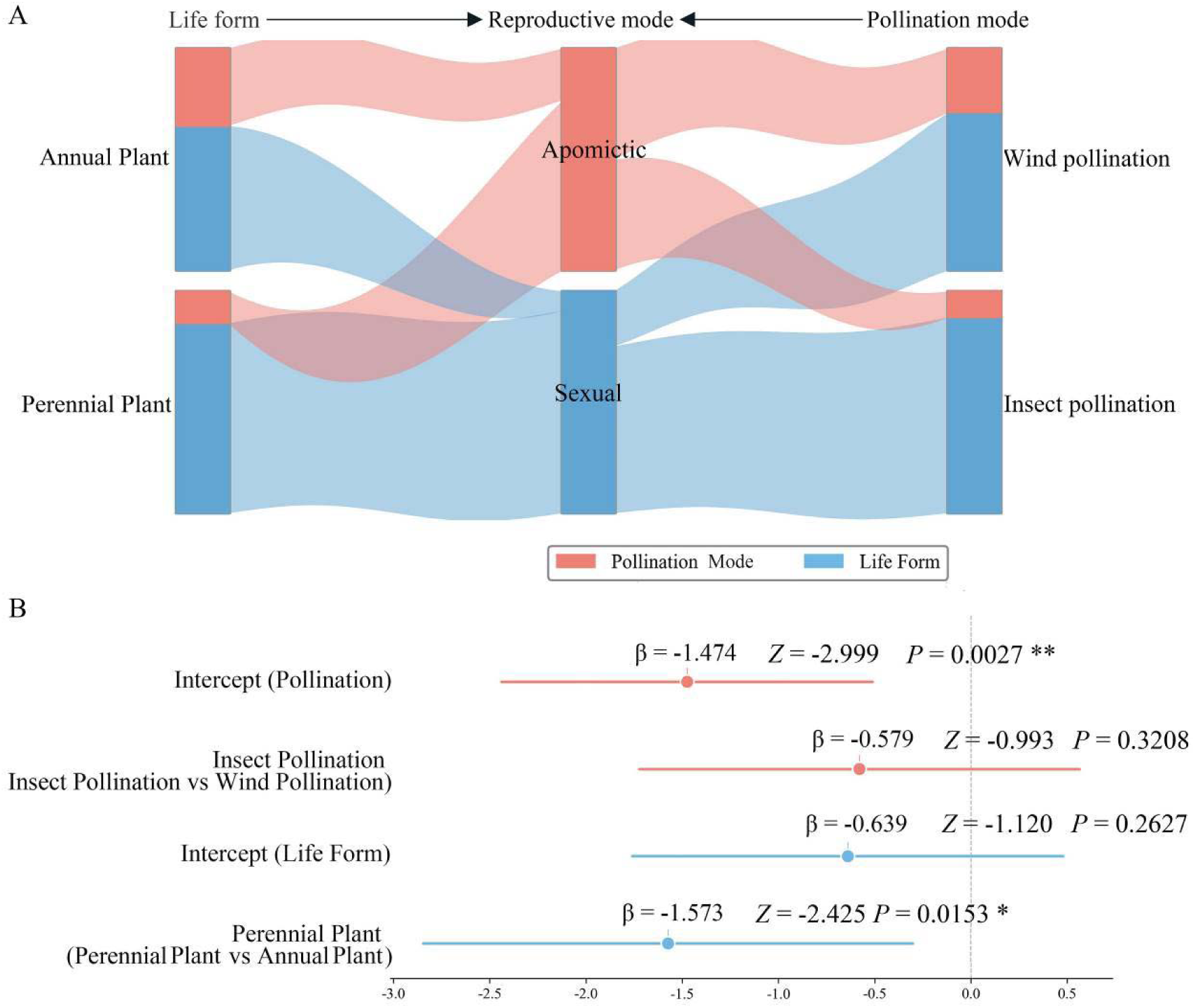
Impact of life form and pollination mode on reproductive modes within karst caves. (A) Sankey diagram visualizing the proportional associations of both life forms and pollination methods with reproductive modes. The top directional arrows indicate the specific relationships, demonstrating the distinct flows from life form to reproductive mode, and from pollination mode to reproductive mode, respectively. (B) Forest plot displaying Phylogenetic Generalized Linear Mixed Model (PGLMM) parameter estimates. The results confirm a significant correlation between life form and reproductive mode (P = 0.015), with annual plants showing a higher prevalence of apomixis. Conversely, pollination mode lacks a significant independent effect (P = 0.321) after accounting for phylogenetic clustering.

When examining the phylogenetic distribution of pollination modes, we found that wind-pollinated species were predominantly concentrated within Urticaceae (38/52, 73.1%), whereas biotic-pollinated species were more evenly distributed across families. Among the 15 apomictic wind-pollinated species, 13 (86.7%) belonged to Urticaceae, specifically the genus Elatostema. This strong phylogenetic clustering complicates the interpretation of pollination mode effects, as the observed pattern may reflect lineage-specific pre-adaptations rather than direct selection against wind pollination per se.

## DISCUSSION

At the heart of the GP hypothesis is Baker’s Law which asserts the evolutionary advantage of autonomous reproduction during the colonization of isolated habitats, which the GP hypothesis links to macro-climatic stress and pollination barriers. We find that in mid-elevation karst caves, it is likely barriers to pollination drive a high frequency of apomixis completely independently of thermal stress or UV radiation. Furthermore, taxonomic composition is dictated by pre-adaptations to deep shade combined with a lineage-specific predisposition for autonomous reproduction, resulting in complete phylogenetic turnover between these contrasting apomixis-rich environments.

### Barriers to pollination, rather than climate stress, select for apomixis in subterranean islands

The GP hypothesis proposes that gametophytic apomixis is linked to high-latitude or alpine environments where conditions favour asexual reproductive assurance (Hörandl et al., 2011; Brožová et al., 2019). Working in a non-alpine, mid-latitude isothermal cave environment, where diurnal fluctuations exceed annual ones, we observed an apomixis frequency of 12.9%, which is statistically comparable to high-latitude or alpine environments in the American Cordillera (12.0%–28.2%; Ptáček et al., 2024) and the Himalayan high-alpine zone (6.0%; Brožová et al., 2019). Since B-FCSS screens pooled seeds where dominant sexual peaks may mask low-frequency signals (Matzk et al., 2000), our 12.9% estimate represents a conservative lower bound, implying a higher true frequency. Our results suggest that even in the absence of clades (e.g., Rosaceae, Asteraceae) that typically dominate the apomictic flora in the Northern Hemisphere, the colonization of cave environments results in frequencies of apomixis comparable to those in extreme alpine environments, suggesting an equivalent intensity of selection that is independent of thermal stress. We found no support for a latitudinal or elevational gradient in apomixis frequency, as would be expected by the GP hypothesis (Vandel, 1928; Hörandl, 2006; Vrijenhoek & Parker, 2009). Rather, our GLMM analyses (Fig. 4) reveal stable apomixis frequencies across elevation (P = 0.503), latitudinal (P = 0.665) and temperature (P = 0.691). We propose that the pressure for reproductive assurance is driven by severe barriers to pollination and light limitation (Li et al., 2006; Monro et al., 2018), both of which exert maximum selective pressure immediately upon colonization. Barriers to pollination arise because caves function as microclimatic islands isolated by surrounding agricultural landscapes, which have different macro-climates and species diversity. Consequently, local insect pollinators are unlikely to possess the physiological or behavioral plasticity required to forage effectively across both cave and the surrounding agricultural landscapes (Gaston 2009). Furthermore, in caves anemophily is severely constrained; interior airflow is markedly reduced compared to surface environments and typically follows diurnal inversion patterns (de Freitas et al. 1982, Badino 2010), limiting steady air currents (Lismonde 2002) necessary for effective pollen dispersal.

To further evaluate the impact of these barriers to pollination, we examined the prevalence of apomixis across different pollination syndromes. While we documented a higher apomixis frequency in wind-pollinated (28.8%) compared to insect-pollinated species (11.8%), this difference was not statistically significant after accounting for phylogenetic structure (β = −0.579, Z = −0.993, P = 0.321) and further research incorporating an appropriate experimental design would be necessary to do so. Despite this phylogenetic constraint on analysing specific pollination systems, the overarching selective pressure of the cave environment remains clear.

Exceptionally low Photosynthetically Active Radiation (PAR) did not significantly impact the frequency of apomixis in karst cave environments; as indicated by the observation that rates between cave entrances and twilight zones were statistically indistinguishable (P = 0.968). This spatial homogeneity suggests that the cave’s combined ecological constraints act as a uniform evolutionary filter, independent of fine-scale variations in PAR. Even at cave entrances, severe pollinator restriction (Li et al., 2006; Wang et al., 2013) favours apomixis as a stable long-term strategy rather than an evolutionary dead-end (Richards, 2003; Hojsgaard & Hörandl, 2015). In fragmented karst landscapes, this autonomous reproduction provides a decisive colonization advantage, suggesting that barriers to pollination actively select for apomixis in a manner consistent with Baker’s Law (Baker, 1967).

### Phylogenetic turnover and life-history constraints: the role of pre-adaptation

Consistent with the hypothesis pollination barriers acting as ecological filters, our results indicate a significant phylogenetic signal (P = 0.002) with genus-level clustering, suggesting that the transition to asexuality in cave environments is phylogenetically constrained. Previous reviews of apomixis across both animals (Murphy et al., 2000; Freitas et al., 2022) and plants (Hojsgaard et al., 2014; Hörandl et al., 2024) have consistently identified phylogenetic constraint as a major determinant of autonomous reproduction. While global temperate hotspots of apomixis are characterized by the dominance of apomictic lineages in the Poaceae, Rosaceae, and Asteraceae (comprising approx. 75% of apomicts; Hojsgaard et al., 2014), our cave ecosystem dataset demonstrates extensive phylogenetic turnover relative to these (Webb et al., 2002; Svenning et al., 2008; Cai et al., 2025). The Poaceae, Rosaceae, and Asteraceae were either absent or exclusively sexual in our samples (Table S2). Instead, the Urticaceae (Elatostema), Euphorbiaceae (Acalypha, Mallotus), and Vitaceae (Causonis) are the dominant apomictic lineages, accounting for 40.9%, 9.1%, and 9.1% of the asexual populations, respectively. This distinct clustering of apomixis within certain lineages suggests that in cave ecosystems, apomixis is a lineage-specific trait dependent on evolutionary pre-conditions rather than a generalized and convergent response to pollination stress. Our two contingency-table analyses support this interpretation, family composition differed among ecosystems whether the analysis was restricted to six focal families or included an Other-families category (both P < 0.001). The robustness of the result to the broader family grouping strengthens the evidence for lineage turnover.

The prevalence of Elatostema in the karst caves illustrates the intersection of both phylogenetic pre-adaptation and ecological filtering. Elatostema exhibits ancestral physiological shade tolerance through its association with the forest understory habitat (Wang, 2014), enabling vegetative persistence under light-limited conditions where typical heliophytic taxa (e.g., Asteraceae) fail to establish. Elatostema also shows a high frequency of polyploidy and hybridization (Fu et al., 2017, 2025), mechanisms associated with apomeiosis (Carman, 1997; Bicknell & Koltunow, 2004; Hojsgaard & Hörandl, 2019). Under the presumed reproductive constraints of the cave environment, selection acts against obligate sexual populations, this genomic plasticity may facilitate the generation of unreduced gametes, followed by the fixation of apomictic genotypes (Vrijenhoek, 1984; Mason & Pires, 2015; Pannell et al., 2015). We propose, therefore, that the combination of ancestral shade tolerance and genomic plasticity provides the basis for colonization and reproductive assurance in the karst cave habitat.

In addition to the phylogenetic turnover, our observations also identify a life-history divergence (Morgan, 2001; Richards, 2003), karst caves exhibiting a higher for apomixis amongst annual species (35.0% compared to perennial taxa (14.5%; β =-1.573, Z =-2.425, P = 0.015), in contrast to high elevation montane habitats where XXX. In contrast, apomixis in temperate and alpine regions is frequently correlated with a perennial life history to ensure long-term persistence (Klimešová et al., 2011; Barrett, 2015), the correlation with annuals in cave environments is initially surprising, as caves are stable, aseasonal habitats comprising several perennial lineages (e.g., Elatostema, Acalypha). The correlation between apomixis and annual life history suggests that single-season reproductive constraints act as a critical evolutionary driver (McNamara et al., 2011; Brannelly et al., 2016). Under this scenario, the selective pressure for immediate reproductive assurance is elevated for annual species seeking to colonise caves, which have a limited temporal window to complete their life cycle. This increases the selective advantage of autonomous reproduction, promoting the fixation of apomixis as a mechanism of reproductive compensation (Krams et al., 2011). Conversely, perennial (iteroparous) taxa can rely on vegetative persistence to buffer against pollinator failure. In annual lineages, therefore, this temporal constraint.

Despite this life-history divergence, the concentration of variance at the genus level (Variance = 1.139–1.259) indicates that the relationship between shade tolerance and reproductive mode remains evolutionarily conserved. The low variance at the family and species levels suggests that following the colonization of the cave habitat, apomixis functions as a stable reproductive strategy constrained primarily at the generic rank. Forest-derived lineages have inhabited East Asian caves since the Oligocene-Miocene boundary (Li et al., 2022); within these isolated environments, severe light limitation and barriers to pollination likely interacted with pre-existing genetic architectures to maintain reproductive output (Carman, 1997; Bicknell & Koltunow, 2004). Accordingly, apomixis in karst caves is not a novel adaptation, but as a phylogenetically constrained more general response to conditions that limit sexual reproduction.

### Reassessing Geographical Parthenogenesis: sampling biases and environmental filtering

The GP hypothesis was originally proposed to explain spatial variation in the frequency of apomixis across landscapes, heavily relying on case studies from extreme, high-elevation climates. However, our findings suggest that the theoretical consensus informing GP may be influenced by methodological constraints and ecologically biased sampling. As summarized in Table 2, recent community-level surveys of apomixis have focused on environments characterized by macro-climatic thermal and UV stress, such as the European Alps, the Himalayas, and the American Cordillera. Consequently, the dominant apomictic lineages identified in these regions are restricted primarily to three families: Asteraceae, Rosaceae, and Poaceae.

By sampling a mid-elevation, isothermal environment characterized by biotic stressors, specifically light limitation and barriers to pollination, we demonstrate that the ecological drivers of apomixis are more diverse than the GP hypothesis accounts for. Despite lacking the thermal extremes of alpine zones, the karst cave flora exhibits an apomixis frequency (12.9%) that is comparable to high-latitude and high-elevation regions. This is accompanied by a phylogenetic turnover, with shade-tolerant lineages such as Urticaceae, Euphorbiaceae, and Vitaceae comprising the asexual assembly (Table 2), which suggests that barriers to pollination act as an independent evolutionary filter. There are many habitats where vascular plant populations can become reproductively isolated, especially amongst wind-pollinated taxa. For example, dense forest understories restrict the aerodynamic currents necessary for pollen transport (Whitehead, 1969; Culley et al., 2002), while topographical barriers and habitat fragmentation dilute pollen concentrations below the thresholds required for effective outcrossing (Knapp et al., 2001; Aguilar et al., 2006). Such structurally and topographically isolated habitats occur across a wide range of elevations and latitudes. Importantly, they represent a much greater proportion of the vegetated terrestrial environment than high-elevation mountains; forests occupy over 30% of the global land area (Scogings, 2023; FAO, 2020), compared to the approximately 3% for the alpine zones (Körner et al., 2011). However, prior to this study, the frequency of apomixis within these geographically extensive but structurally isolated environments had not been systematically evaluated (see Table 2). To further test the role of reproductive isolation, future studies should evaluate the frequency of apomixis in dense forests and steeply dissected landscapes across tropical and subtropical latitudes

In addition, FCSS, which has emerged as the high-throughput standard for characterizing reproductive pathways in biodiversity hotspots (Matzk et al., 2000; Hörandl et al., 2011; Brožová et al., 2019; Ptáček et al., 2024), uncovering macro-evolutionary patterns that traditional embryology might obscure (Schwendener, 1969; Czapik, 1975). It can, however, also introduce a source of bias.as B-FCSS relies on detectable endosperm signals, a requirement that can introduce systematic bias by excluding non-endospermic taxa (Matzk et al., 2000; Matzk, 2007; Ptáček et al., 2024). This limitation notably affected our analysis of Gesneriaceae, a dominant family in cave floras (Fu et al., 2022) that typically produces seeds devoid of endosperm (Li & Wang, 2004); consequently, we were able to screen only five of the 118 recorded species. Nevertheless, our sensitivity analysis confirms that even if all unscreened Gesneriaceae are treated as sexual, the apomixis frequency in cave floras (7.7%) still exceeds the global average (ca. 2.2%; Hojsgaard et al., 2014) and that of regional high-mountain floras (4.7%–6.0%; Hörandl et al., 2011; Brožová et al., 2019). While higher frequencies (12.0%–28.2%) have been reported in American alpine floras (Ptáček et al., 2024), those estimates are likely also inflated by the exclusion of non-endospermic families such as Fabaceae and Orchidaceae, as noted by the authors. This highlights how methodological limitations in broad-scale screenings can skew our perception of where apomixis most frequently occurs.

Interpretive challenges also arise when the embryo and endosperm possess identical ploidy levels (Williams & Friedman, 2002; Kao, 2007), rendering DNA ratios uninformative. A specific technical difficulty exists in distinguishing autonomous endosperm (4C) from the embryo G2 phase (Pind = 2.0). While some studies omit G2 peaks (Šarhanová et al., 2012; Brožová et al., 2019; Lepší et al., 2019) or prioritize endosperm signals (Hörandl et al., 2011; Dobeš et al., 2013), we adopted a rigorous comparative approach, validating a 4C peak as authentic endosperm only if its nuclei count ratio (Rsample) reached at least 75% of the baseline from related sexual species (Rref). This threshold effectively filters background cell-cycle noise, ensuring that our 12.9% frequency estimate reflects true reproductive transitions rather than an overestimation driven by G2-phase artifacts.

Ultimately, these results demonstrate that when methodological biases are rigorously controlled for, the evolutionary transition to asexuality is revealed to be far more taxonomically diverse and ecologically decoupled from macro-climates than the GP hypothesis traditionally suggests.

### Conservation Implications for Karst Endemics

The prevalence of apomixis in karst caves necessitates a shift from broad climatic conservation models to strategies treating individual caves as discrete evolutionary units. Because apomixis stabilizes highly adapted genotypes within fragmented landscapes, maintaining the specific microenvironments of individual caves is critical. Anthropogenic alterations to light or humidity regimes at cave entrances may disrupt the selective filters that sustain these specialized asexual lineages (Ren et al., 2021).

Furthermore, the observed phylogenetic turnover implies that conservation management must be tailored to specific evolutionary lineages. For dominant apomicts like Elatostema, priority should be given to protecting polyploid apomictic complexes that function as vital genetic reservoirs in resource-poor habitats (Fu et al., 2025). These populations also serve as invaluable germplasm repositories, offering fixed, adapted genotypes essential for future karst forest restoration (Clements et al., 2006; Monro et al., 2018). Recognizing these subterranean habitats as dynamic sites of reproductive innovation, rather than mere physical refugia, is fundamental to ensuring the long-term evolutionary resilience of the unique karst flora (Li et al., 2022).

## CONCLUSION

This study shows that gametophytic apomixis occurs in subtropical karst caves at a frequency of 12.9%, comparable to global high-latitude hotspots. Apomixis frequency remains statistically stable across elevational and latitudinal gradients, consistent with a spatial decoupling from macro-climatic thermal stress. This pattern suggests that micro-environmental constraints in caves, including barriers to pollination and light restriction, are associated with reproductive assurance strategies independent of geographic position. The cave environment is further associated with a phylogenetic turnover in which apomixis is concentrated in shade-tolerant lineages such as Urticaceae, Euphorbiaceae, and Vitaceae, rather than in traditional asexual-prone families such as Asteraceae and Poaceae.

These findings indicate that the evolutionary transition to asexuality is more taxonomically diverse than previously recognized. Apomixis is characterized as a threshold response to subterranean colonization rather than a linear reaction to climatic variation. The association between apomixis and annuality is consistent with single-season reproductive constraints acting as a life-history bottleneck that favours semelparous lineages capable of autonomous seed set. Morphologically, the predominance of female inflorescences in apomictic Elatostema populations is consistent with reduced investment in male function under limited pollinator availability. These results support treating individual caves as discrete evolutionary units where reproductive strategies are decoupled from surface environmental gradients. The occurrence of apomixis in non-model lineages such as Elatostema provides a framework for investigating the genetic basis of asexual reproduction in fragmented and extreme environments.

## RESOURCE AVAILABILITY

### Lead contact

Further information and requests should be directed to the lead contact, Yi-Gang Wei.

### Materials availability

This study did not generate new, unique reagents.

### Data and code availability

Data supporting the findings are provided in the article and supplemental tables. Analysis code is available from the lead contact upon reasonable request and should be deposited in a public repository before acceptance.

## ACKNOWLEDGMENTS

xxx

## STAR METHODS

Study Area and Plant Sampling

Field sampling was conducted across 67 subtropical karst cave systems in Southwest China, a global biodiversity hotspot for cave-dwelling flora (Monro et al., 2018). To ensure broad taxonomic and ecological representation, we investigated 305 populations comprising 203 angiosperm species. Sampling was strictly confined to the entrance and twilight zones, which are defined by Monro et al. (2018) as the limits of photosynthetically active radiation capable of supporting vascular plants. The dark zone, being devoid of autotrophic vascular flora, was excluded from this study. Detailed geographical coordinates, light zone classifications, and population counts for each site are consolidated in Table S1–S2.

To capture the representative reproductive characteristics of each population, mature seeds were harvested from multiple individuals (typically > 10 whenever available) to construct a comprehensive seed pool for each accession. These seeds were stored in labeled paper bags and air-dried under ambient conditions to maintain stability before analysis. For taxonomic verification, herbarium voucher specimens were collected for every population. Identifications were performed by consulting relevant taxonomic literature on cave flora (Fu et al., 2022; Wei et al., 2025) and online databases such as the Chinese Virtual Herbarium (CVH; https://www.cvh.ac.cn/) and the Plant Photo Bank of China (PPBC; https://ppbc.iplant.cn/), and deposited at the Herbarium of Guangxi Institute of Botany (IBK).

### Reproductive Mode Screening via Bulked Flow Cytometry (B-FCSS)

The reproductive mode of each sampled population was determined using Bulked Flow Cytometric Seed Screen (B-FCSS), an optimized high-throughput approach specifically developed for large-scale population surveys (Matzk et al., 2000; Dobeš et al., 2013). By pooling more than 10 mature seeds per sample, we obtained a representative population-level signature of the reproductive pathway, a strategy widely encouraged for screening broad taxonomic and ecological representation while maintaining statistical robustness (Hörandl et al., 2008; Kolarčik et al., 2018).

The flow cytometry protocol was optimized for a wide range of cave-dwelling species. After removing any persistent fruit tissue or seed coats, the seeds were placed in a petri dish with 0.5 mL of ice-cold Otto I lysis buffer (0.1 M citric acid, 0.5% v/v Tween 20) and finely minced with a sharp razor blade for approximately 60 seconds to release the nuclei. The resulting suspension was filtered through a 42-μm nylon mesh. Subsequently, 1 mL of Otto II staining buffer (0.4 M Na2HPO4·12H2O) containing the DNA-intercalating fluorochrome Propidium Iodide (PI) was added. Samples were incubated in the dark for 10–20 minutes at room temperature to ensure complete staining. Analyses were performed on a Partec CyFlow ML flow cytometer. For each sample, sufficient particles (typically > 3,000) were collected to ensure the generation of high-resolution histograms and reliable peak identification.

### Inference and Classification Framework

The reproductive pathway was inferred based on the peak index (Pind), defined as the ratio of the mean fluorescence intensities of the endosperm to embryo nuclei (Dobeš et al., 2013). Following the framework of Schinkel et al. (2016) and Kolarčik et al. (2018), we classified reproductive modes as follows: sexual (Pind ≈ 1.5), autonomous apomixis (Pind ≈ 2.0), and pseudogamous apomixis (Pind ≈ 2.5 or 3.0). Seeds exhibiting atypical ratios between 1.6 and 1.85 were classified as having an Intermediate Reproductive Mode (IRM). While this range aligns with the definition of hemisexuality described in Rosa (Kolarčik et al., 2018), mathematically it is also indistinguishable from maternal BIII hybridization (fertilization of unreduced egg cells, Pind ≈ 1.67; Hojsgaard & Hörandl, 2019). Seeds exhibiting atypical ratios between 1.1 and 1.4 were classified as sex (paternal BIII hybridization). Given the bulked nature of our sampling strategy, the detection of a dominant peak in this range indicates a stabilized, population-level strategy rather than sporadic fertilization events.

To rigorously address biological ambiguity while maintaining statistical comparability with global datasets, we implemented a Dual-Classification Framework:

- *Index A (Standardized Apomixis Frequency - Strict Definition)*: This index includes only populations with definitive autonomous (Pind ≈ 2.0) or pseudogamous (Pind ≥ 2.5 or 3.0) signals. Intermediate forms (1.6–1.85) are treated as non-apomictic or excluded. This conservative metric ensures strict comparability with previous large-scale screenings in the Alps and Andes (e.g., Ptáček et al., 2024).
- *Index B (Broad-sense Apomictic Capacity - Functional Definition)*: This index groups Intermediate Reproductive Modes (1.6–1.85) with apomictic lineages. This classification reflects the functional capacity for apomeiosis (formation of unreduced gametes), representing a ‘reproductive assurance’ mechanism where plants deviate from strict sexual reproduction, regardless of whether the unreduced egg is subsequently fertilized (BIII) or develops parthenogenetically.

To distinguish functional autonomous endosperm (4C) from G2-phase embryo nuclei artifacts (4C), we implemented a strict dual-validation strategy rooted in seed stoichiometry. 4C signals were classified as autonomous endosperm only if they exceeded an 8% abundance threshold relative to the 2C peak. For accessions exhibiting a lower frequency 4C peak, a 4C peak was classified as authentic endosperm only if its nuclei count ratio (Rsample) was comparable in magnitude to a taxonomic baseline (Rref) established from phylogenetically related sexual species (typically Rsample ≥ 75% × Rref); otherwise, it was categorized as background cell cycle noise. Accessions failing these criteria were classified as ‘na’ and excluded.

### Testing Hypothesis 1: Apomixis Frequency and Regional Comparisons

To place our findings in a global context and test whether apomixis occurs at a higher rate in caves, the frequency of apomixis in the subtropical karst cave flora was compared to frequencies reported from the American Cordillera (Ptáček et al., 2024) and the Himalayas (Brožová et al., 2019). We utilized Index A (Standardized Apomixis Frequency) for all inter-regional comparisons to ensure methodological consistency. We performed z-tests for the difference of two independent proportions to compare prevalence across regions and distinct light zones (entrance vs. twilight). For comparisons involving small sample sizes (N < 40), Conover’s continuity correction was applied to maintain statistical robustness (Martin Andres et al., 2024). To address potential bias from excluding non-endospermic taxa—specifically the highly diverse Gesneriaceae, which typically lack a developed endosperm (Li & Wang, 2004) rendering B-FCSS ineffective—we incorporated a conservative sensitivity analysis. Because cytological evidence confirms the vast majority of Primulina are stable diploids (Kang et al., 2014) and apomixis is rare in diploids (Hojsgaard et al., 2014), we treated these excluded taxa as sexually reproducing to validate the lower bounds of our estimated apomixis frequencies. All statistical analyses were performed in R (v.4.1.1; R Core Team, 2021).

### Testing Hypotheses 2 & 3: Phylogenetic Signal and Lineage Pre-adaptation

To assess whether cave apomicts occur in lineages associated with the GP hypothesis or are drawn from the local forest understory, a phylogenetic tree of the sampled species was constructed using the ‘V.PhyloMaker’ R package (v.0.1.0; Jin & Qian, 2019). We applied scenario 3, which generates phylogenies by pruning a comprehensive mega-tree of vascular plants (GBOTB.extended). Consistent with previous large-scale surveys of alpine floras (Ptáček et al., 2024), this analysis was restricted to species exhibiting either sexual or apomictic reproduction to maintain binary contrast. The phylogenetic signal for apomixis (coded as 0 for sexual and 1 for apomictic) was tested using phylogenetic logistic regression via the R package ‘ape’ (v.5.5; Paradis et al., 2004). A significant signal indicates that reproductive modes are phylogenetically conserved due to common ancestry, while a non-significant signal suggests independent evolution across lineages. The tree was visualized using TvBOT (v2.6.1) (Xie et al., 2023).

To test compositional turnover directly, we tabulated unique apomictic species by family across five ecosystems. A 5 x 6 table included Urticaceae, Euphorbiaceae, Vitaceae, Asteraceae, Poaceae, and Rosaceae; a second 5 x 7 table added all remaining families as an Other category. Because expected counts were sparse, both tables were evaluated using two-sided Fisher-Freeman-Halton exact tests with Monte Carlo P values (10,000 replicates; set.seed(123)).

### Evaluating Drivers: Decoupling Macro-Climates from Micro-Habitats

To analyze the influence of environmental and biological factors, we employed Generalized Linear Mixed Models (GLMMs) with a binomial error distribution and logit link function (Bolker et al., 2009). For these ecological models, we utilized Index B (Broad-sense Apomictic Capacity) as the response variable, positing that the functional ability to produce unreduced gametes represents the critical adaptive trait for reproductive assurance in resource-limited cave environments. The analytical dataset was subsetted to include 261 populations (174 species). Given the distinct ecological scales of our predictors, we constructed separate models. Environmental drivers (altitude and latitude) were modeled at the population level to test for spatial decoupling from macro-climatic gradients, while biological traits (life form and pollination mode) were modeled at the species level. To account for taxonomic non-independence, family, genus, and species were incorporated as nested random effects, allowing us to partition the variance across different evolutionary ranks (Zuur et al., 2009). Pollination mode was a documented species-level syndrome. We did not measure pollinator visitation, pollen deposition, cave airflow, or experimental pollen limitation; the analysis therefore tests association with pollination mode rather than a direct effect of pollination barriers.

## AUTHOR CONTRIBUTIONS

[Authors should insert a CRediT contribution statement before submission.]

## DECLARATION of GENERATIVE AI AND AI-ASSISTED TECHNOLOGIES IN THE WRITING PROCESS

During preparation of this work, the authors used OpenAI Codex and Gemini 3.1 Pro to improve readability, language, and manuscript organization. The authors reviewed and edited the content and take full responsibility for the content of the article.

## SUPPLEMENTAL INFORMATION

Document S1. Supplemental Tables S1 and S2.

